# Hepatic Xanthine Oxidoreductase Sustains Antithrombotic Nitric Oxide Signalling Through the Nitrate–Nitrite-Nitric Oxide Pathway

**DOI:** 10.64898/2026.09.13.751308

**Authors:** Tipparat Parakaw, Cristina Perez Ternero, Federica Filomena, Nicki Dyson, Harriet Allan, Najada Cufaj, Gianmichele Massimo, Mike Curtis, Rayomand S Khambata, Amrita Ahluwalia

## Abstract

**Background:** Tonic endothelium-derived nitric oxide (NO) suppresses platelet activation physiologically, and its loss underlies the thrombotic risk of endothelial dysfunction. Inorganic nitrate and nitrite provide an alternative NO source, and xanthine oxidoreductase (XOR) is a candidate nitrite reductase, but whether endogenous XOR sustains platelet NO signalling *in vivo*, and its source is unknown.

**Methods:** Dietary nitrate (15 mmol/L KNO₃) was given to *eNOS*^−/−^ and *ApoE*^−/−^ mice. XOR was interrogated pharmacologically (allopurinol) and genetically (global *Xdh*^+/−^ and hepatocyte-specific XOR knockout, HXOR KO). Haemostasis and thrombosis were assessed by tail bleeding and intravital microscopy imaging of FeCl₃-induced mesenteric arterial thrombosis, alongside aggregometry, flow cytometry, platelet VASP^Ser239^ phosphorylation, cGMP and ozone chemiluminescence-based analysis of nitrate and nitrite.

**Results:** Dietary nitrate raised plasma nitrate and nitrite levels in all genotypes. In *eNOS*^−/−^ and *ApoE*^−/−^ mice, it prolonged bleeding time and normalised thrombus burden. In *ApoE*^−/−^ mice nitrate treatment improved vasorelaxation, without altering XOR or eNOS expression. Allopurinol suppressed nitrite reductase activity in liver and plasma but not aorta, elevated plasma nitrite, shortened bleeding time and reduced platelet P-VASP^Ser239^ expression. *Xdh*^+/−^ mice were spontaneously prothrombotic, showed reduced P-VASP^Ser239^ and exaggerated calcium mobilisation, and were refractory to dietary nitrate treatment despite equivalent nitrite elevation. XOR was undetectable in platelets. HXOR KO mice phenocopied global deficiency, with blunted nitrite-induced vasorelaxation but preserved acetylcholine and spermine-NO responses, reduced P-VASP^Ser239^, enhanced aggregation and shortened bleeding time.

**Conclusions:** Hepatic XOR sustains platelet NO-cGMP signalling and thromboresistance through an inter-organ, endocrine-like axis. Dietary nitrate restores antithrombotic protection when endothelial NO generation fails, whereas XOR inhibition removes physiological tonic antithrombotic signal.

## INTRODUCTION

The vascular endothelium is a critical regulator of cardiovascular homeostasis through the continuous generation of nitric oxide (NO)^1^. Beyond its vasodilator actions, endothelial-derived NO suppresses platelet activation, adhesion and aggregation through activation of soluble guanylyl cyclase (sGC), increased cyclic guanosine monophosphate (cGMP) production and downstream protein kinase G signalling in platelets^2–4^. This pathway limits intracellular calcium mobilisation, inhibits granule secretion and prevents activation of integrin αIIbβ3, thereby maintaining blood fluidity and protecting against thrombosis^5–8^. Conversely, impaired NO bioavailability is a hallmark of endothelial dysfunction and contributes to enhanced platelet reactivity and thrombotic risk in disorders including atherosclerosis, hypertension and diabetes^9^. Thus, the concept that restoring endothelial NO levels might be a useful strategy in limiting disease progression has been much considered. In this context, a promising approach to improve NO levels is through the use of inorganic nitrite (NO₂⁻) or nitrate (NO_3_⁻).

In addition to the canonical L-arginine-NO synthase pathway, NO can be generated through sequential reduction of inorganic nitrate and nitrite. This pathway provides an alternative source of bioactive NO that can be augmented through the consumption of NO_3_⁻-rich vegetables or through NO_3_⁻ salt ingestion^10^. Xanthine oxidoreductase (XOR) is a major mammalian nitrite reductase and is upregulated in several cardiovascular diseases, raising the possibility that increased XOR activity may enhance nitrate-derived NO generation under pathological conditions^11,12^.

It has been shown that dietary NO_3_⁻ or NO_3_⁻ salt supplementation attenuates *ex vivo* assessed platelet reactivity in humans and experimental models^13–15^, but the mechanisms responsible and the contribution of endogenous XOR remain incompletely understood. Although traditionally viewed as a source of oxidative imbalance XOR has also been identified as a physiologically important nitrite reductase generating NO^16^. However, whether this pathway contributes to platelet homeostasis *in vi*vo, and might be exploited to restore antithrombotic signalling during endothelial dysfunction, remains unknown. Moreover, the tissue source of functionally relevant XOR-derived NO has not been defined.

We therefore tested the hypothesis that XOR-derived NO restrains platelet activation and thrombosis through cGMP-dependent signalling and that dietary nitrate restores platelet function in states of endothelial dysfunction by enhancing this pathway. Using eNOS knockout (*eNOS*^-/-^)^17^ and the atherosclerotic *ApoE*^-/-^ mouse^18^, together with global and hepatocyte-specific XOR-deficient models,^16,19^ we identify hepatic XOR as a critical regulator of nitrate-derived NO signalling and platelet homeostasis *in vivo*.

## Methods

All reagents were purchased from Sigma Chemical Co. unless otherwise stated. See supplement for full experimental details.

### Animals and dietary treatments

All animal experiments were conducted in accordance with the Animals (Scientific Procedures) Act 1986, UK and had the approval of the UK Home Office and Local Animal Welfare Ethical Review Board Committee.

Constitutive global XOR-deficient (*Xdh^+/-^* and *Xdh^-/-^*)^19^ and hepatocyte-specific XOR-knockout (HXOR KO)^16^ mice together with their littermate controls, *Xdh*^+/+^ and *Xdh^fl/fl^* (WT) were generated in-house as described previously. In some experiments, constitutive global apolipoprotein E (*ApoE*^-/-^) and endothelial nitric oxide synthase (*eNOS*^-/-^)^20^ knockout mice from our in-house-maintained colonies were used. All studies were sex-balanced and performed on 4-8 week-old littermate wild-type and mutant animals.

For the NO_3_^-^ treatment studies, mice were randomly assigned (using an online randomisation code generator) to control (15 mM KCl) or treatment (15 mM KNO_3_) groups. Supplementation was provided in the drinking water, which was replaced every 2-3 days. This dosing regimen was chosen based on our previous work showing that 15 mM KNO_3_ in the drinking water produces a robust elevation in plasma NO_3_^-^ and NO_2_^-^ levels^2, 3^, with the latter comparable to those levels observed in humans. Similarly, for allopurinol treatment, mice received 1 mM allopurinol (in 1.2 mM NaOH) or vehicle (1.2 mM NaOH) in the drinking water for two weeks, with supplemented water replaced every 2-3 days as previously described^21^.

### Blood collection and platelet isolation

Blood was collected by cardiac puncture using 3.8% (w/v) sodium citrate as anticoagulant. Fresh blood was used for impedance aggregometry or flow cytometry analysis (see below). A second aliquot was centrifuged at 13,000 g for 5 minutes at 4°C to obtain plasma, which was snap-frozen and stored at −80°C for NO_x_ analysis. A third aliquot was used for platelet isolation for cGMP quantification and Western blot analyses.

### Tail bleeding assay

A modified tail bleeding assay was used to assess *in vivo* platelet function^22^. Briefly, animals were anaesthetised with isoflurane (IsoFlo, Zoetis, UK), and body temperature was maintained at 37°C throughout the procedure. Bleeding was initiated by excising 0.25 cm of the distal tip of the tail using a scalpel blade, after which the tail was immediately immersed in warm saline (37°C), and the time to cessation of bleeding was recorded. For the NO_3_^-^ treatment study, tail excision length was adjusted based on body weight: 0.5 cm in mice >20 g and 0.4 cm in mice <20 g. A priori exclusion criterion was applied whereby any animal with a bleeding time exceeding 10 minutes was excluded from analysis.

### FeCl_3_-induced arterial thrombosis

To simulate endothelial damage-triggered thrombosis FeCl_3_-induced chemical damage was used^23^. Four-week-old littermate mice were anesthetised with a cocktail of ketamine (Ketaset, 150 mg/kg body weight, Pfizer) and xylazine (Rompun, 7.5 mg/kg body weight, Bayer AG) and body temperature was maintained at 37°C throughout the procedure. The platelet-specific anti-Glycoprotein Ib (CD42) antibody conjugated with DyLight488 (0.1 μg/g body weight, Emfret Analytics, RRID: AB2890921) was administered via the tail vein^24^. The mesentery was exteriorised under an inverted microscope and kept hydrated by superfusion with 5% CO_2_/95% N_2_-gassed bicarbonate buffer saline (0.13 M NaCl, 3 mM KCl, 1 mM MgSO_4_, 18 mM NaHCO_3_, 2 mM CaCl_2_). A 1 x 3 mm strip of Whatman filter paper soaked in 10% FeCl_3_ was topically applied to a mesenteric artery (100-150 μm in diameter) for 1 minute and rinsed with bicarbonate buffer saline. Fluorescent images were recorded every 5 seconds for 45 minutes using a Zeiss Axioskop microscope (20x objective) and VideoVelocity Time-Lapse (Candilabs) software. Three images per time point were analysed using ImageJ (NIH) to determine total arterial area and DyLight488-positive thrombus area at 0, 5, 10, 15, 20, 25, 30, 35, 40, 45 minutes.

### Vascular pharmacology

Vascular reactivity of aortic ring preparations was assessed using classical tissue bath pharmacology.

### Impedance aggregometry: *ex vivo* platelet aggregation responses

Impedance aggregometry was used to assess *ex vivo* platelet aggregation in response to a range of agonists using a MultiplateR analyzer (Dyabyte Medical, Germany).

### Flow cytometry

Flow cytometry analysis was used to determine platelet count and size, platelet P-selectin expression and calcium flux under basal conditions and after agonist stimulation, before and after agonist-induced activation, platelet calcium flux in response to agonist activation, platelet-monocyte aggregates or platelet-neutrophil aggregates, and intracellular and extracellular platelet XOR expression. All analyses were conducted on a BD LSR Fortessa cell analyser (BD Biosciences, UK).

### Ozone chemiluminescence

Plasma, aorta and liver NO_3_^-^ and NO_2_^-^ concentrations were determined by ozone-based chemiluminescence using a Sievers nitric oxide analyser (NOA 280i, Analytix, Tyne and Wear, UK) as previously described elsewhere^25,26^.

### Nitrite reductase activity

Nitrite reductase activity was evaluated in plasma, thoracic aorta and liver homogenates using a Sievers nitric oxide analyser (NOA 280i, Analytix, Tyne and Wear, UK) as previously described^11,27^.

### Western blot analysis

XOR, eNOS and VASP expression were assessed in platelets, aorta and liver using Western blotting and selective antibodies, expression visualised using enhanced chemiluminescence (Bio-Rad, CA, USA) and densitometric analyses conducted using ImageJ software.

### cGMP ELISA of platelet pellets

Platelets were isolated as described above, stored at −80°C and lysed on the day of the assay with a 0.5% solution of dodecyl trimethylammonium bromide. cGMP levels were determined by an enzyme immunoassay kit (Cyclic GMP EIA Kit RPN226, GE Healthcare, Chicago, USA) according to the manufacturer’s protocol for acetylated samples and normalised to protein concentration.

### Statistical analysis

All estimated effect sizes were calculated using our in-house preliminary data (see supplement for full details). Values shown are mean ± SEM of n values where n refers to the number of animals. For two group comparison unpaired Students t test was used. For multiple group comparisons two-way ANOVA was used followed by Sidak’s multiple post-test. All data was assessed for distribution prior to analysis. For data that did not follow a normal distribution non-parametric statistics were conducted using the Mann Whitney for multiple group comparisons and Kruskal Wallis for post-tests. In all cases post-tests were conducted only if F achieved a P>0.05 and there was no significant variance in homogeneity. In all cases a P<0.05 was considered statistically significant.

## Results

### Dietary nitrate augments systemic NO metabolite availability independent of changes in vascular XOR abundance

Dietary nitrate supplementation statistically significantly increased circulating nitrate and nitrite in *eNOS^+/^*^+^ (WT) and *eNOS^-/-^* mice (**Figure 1A&B**), *ApoE^+/+^* (WT) and *ApoE^-/-^* mice (**Figure 2A&B**) and *Xdh-*deficient mice and their respective wild-type littermates (**Figure 5A&B**). These findings confirm effective activation of the nitrate-nitrite pathway across all experimental models. However, while tissue increases in nitrate were consistent across genotypes (**Supplementary Figure 1A, 1C, 2A, 2C, 3A**), tissue nitrite levels showed greater variability and did not reach statistical significance (**Supplementary Figure 1B, 1D, 2B, 2D and 3B**). Nitrate supplementation did not alter hepatic or vascular XOR expression (**Supplementary Figure 1E-H and 2E-H**), aortic eNOS expression or eNOS Ser1177 phosphorylation (**Supplementary Figure 2I-L and 3C-F**). However, nitrate treatment increased total VASP expression in WT animals, an effect absent in mutant mice (**Supplementary Figure 1I-L, 2M-P and 3G-J**). Platelet count was unaffected by either genotype or nitrate supplementation (**Supplementary Figure 4A and 5A**). Together, these findings indicate that dietary nitrate increases systemic NO metabolite availability without altering the major enzymatic sources of NO generation.

**Figure 1:**
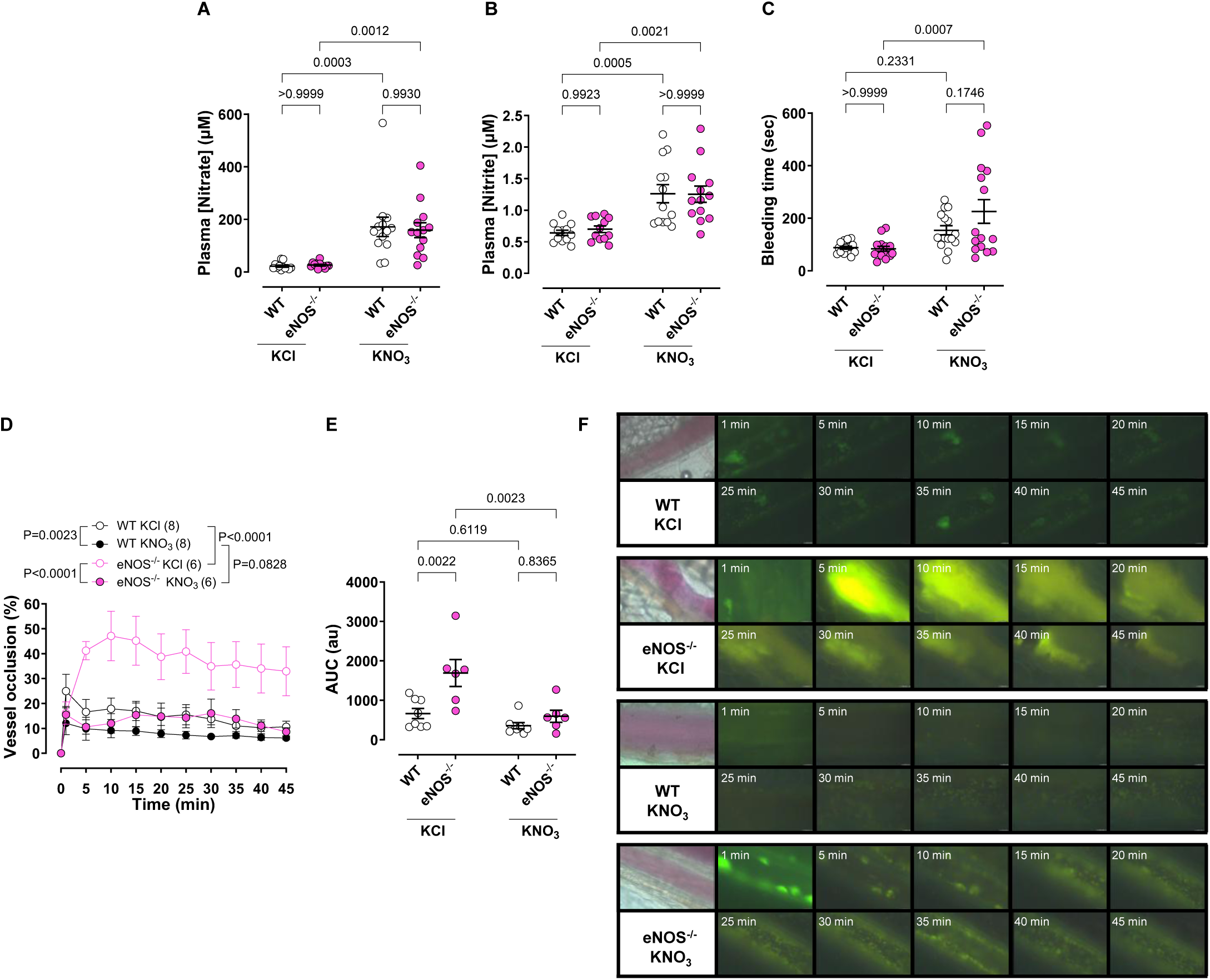
Dietary nitrate reverses the prothrombotic phenotype in eNOS^-/-^ mice. (**A**) Plasma nitrate and (**B**) nitrite concentrations in wild-type (WT) or endothelial nitric oxide synthase-deficient (eNOS^-/-^) mice treated for two weeks with potassium nitrate (15 mM KNO_3_) in the drinking water or potassium chloride (15 mM KCl) as a control. (**C**) Tail bleeding time. (**D**) Vessel occlusion over time, represented as a percentage of thrombus formation to the total vessel area. (**E**) Area under the curve (AUC) of vessel occlusion over time. (**F**) Representative images of thrombus formation. Data are shown as mean ± SEM (shown on the individual graphs). Statistical significance was determined using two-way ANOVA followed by Šídák’s multiple comparisons test.

**Figure 2:**
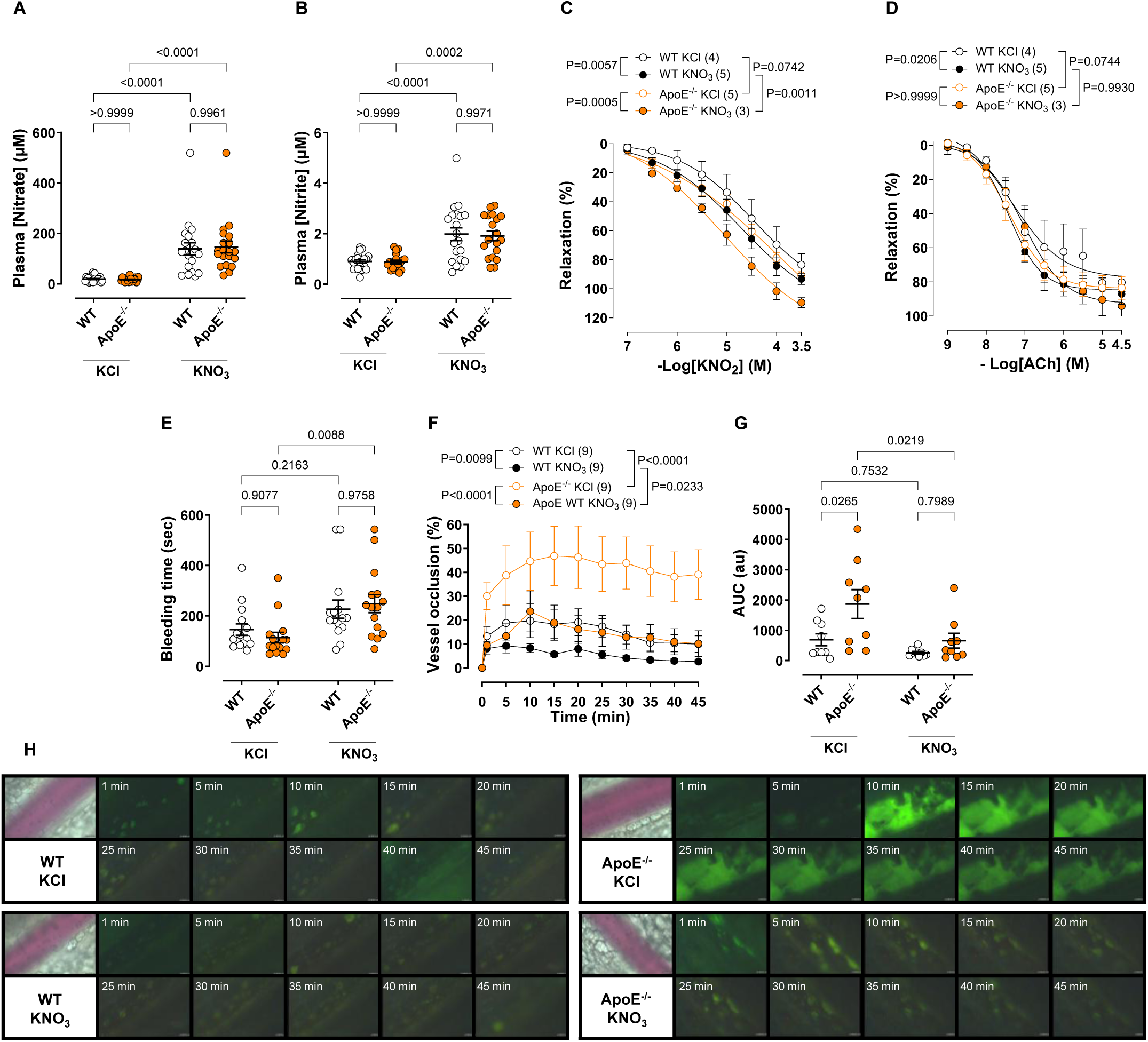
Dietary nitrate improves vascular function and reverses the prothrombotic phenotype in ApoE^-/-^ mice. (**A**) Plasma nitrate and (**B**) nitrite concentrations. (**C**) Potassium nitrite (KNO_2_) and (**D**) acetylcholine (ACh)-induced relaxation of aortic rings precontracted with phenylephrine in wild-type (WT) and apolipoprotein E-deficient (ApoE^-/-^) mice treated for two weeks with potassium nitrate (15 mM KNO_3_) in the drinking water or potassium chloride (15 mM KCl) as a control. (**E**) Tail bleeding time. (**F**) Vessel occlusion over time, represented as a percentage of thrombus formation to the total vessel area. (**G**) Area under the curve (AUC) of vessel occlusion over time. (**H**) Representative images of thrombus formation. Data are shown as mean ± SEM (shown on the individual graphs). Statistical significance was determined using two-way ANOVA followed by Šídák’s multiple comparisons test.

### Dietary nitrate restores antithrombotic function during endothelial dysfunction

Because endothelial NO deficiency promotes platelet hyperreactivity and thrombosis, we first examined whether dietary nitrate could compensate for NO deficiency in *eNOS^-/-^* mice. Platelet P-selectin expression, platelet-monocyte aggregate formation and PAR4-induced platelet aggregation were unaffected by genotype (**Supplementary Figure 4B-E**). Nitrate supplementation had only modest effects on some conventional markers of platelet activation, reducing collagen-induced platelet aggregation *ex vivo*, with a similar trend observed following ADP stimulation (**Supplementary Figure 4F & G**). However, the effects *in vivo* were striking. eNOS^-/-^ mice exhibited enhanced thrombus formation following FeCl₃ injury compared with WT littermates, despite similar bleeding times (**Figure 1C-F**). Dietary nitrate prolonged bleeding time and normalised thrombus burden, effectively restoring haemostatic responses to WT levels (**Figure 1C-F; Supplementary Figure 4**). Importantly, nitrate did not affect thrombus stability or formation kinetics (**Supplementary Figure 4H&I**), suggesting a selective effect on the magnitude of response rather than the dynamics of thrombus formation. These findings establish that dietary nitrate-derived NO can compensate for loss of endothelial NO signalling and restore antithrombotic protection *in vivo*.

A similar protective effect was observed in ApoE^-/-^ mice. Consistent with endothelial dysfunction *ApoE*^-/-^ mice displayed impaired NO responsiveness, characterised by a reduced response to the NO donor spermine-NO (Sper-NO) and augmented contraction to phenylephrine (**Supplementary Figure 5B&C**). Nitrate supplementation improved vasorelaxation to KNO_2_, ACh and Sper-NO (**Figure 2C&D and Supplementary Figure 5B**) while normalising phenylephrine-induced contraction (**Supplementary Figure 5C**). Although *ex vivo* platelet activation remained largely unchanged (**Supplementary Figure 5D-5I**), nitrate supplementation prolonged bleeding time (**Figure 2E**) and reduced thrombus formation to levels observed in WT littermate controls (**Figure 2F-H; Supplementary Figure 5J-L**). Overall, these data demonstrate that dietary nitrate supplementation restores anti-thrombotic function across distinct models of vascular NO deficiency.

### XOR-dependent nitrite-reduction sustains platelet NO-cGMP signalling

To determine whether XOR mediates these protective effects, nitrite reductase activity was assessed in plasma, liver, and aortic homogenates. Nitrite was converted to NO in a concentration-dependent manner in plasma, liver and aortic homogenates (**Figure 3A-C**). Allopurinol significantly suppressed this activity in liver and plasma, although not the aorta. Consistent with inhibition of nitrite reduction *in vivo*, chronic allopurinol treatment increased circulating nitrite concentrations without affecting nitrate levels (**Figure 3D&E**), or nitrite or nitrate levels in the liver (**Supplementary Figure 6A&B**). Loss of XOR activity was accompanied by shortened bleeding time (**Figure 3F**) and reduced phosphorylation of the cGMP-responsive target VASP at Ser239 (**Figure 3G-J**). Together, these findings identify XOR as an important nitrite reductase that sustains platelet NO-cGMP signalling and limits platelet reactivity *in vivo*.

**Figure 3:**
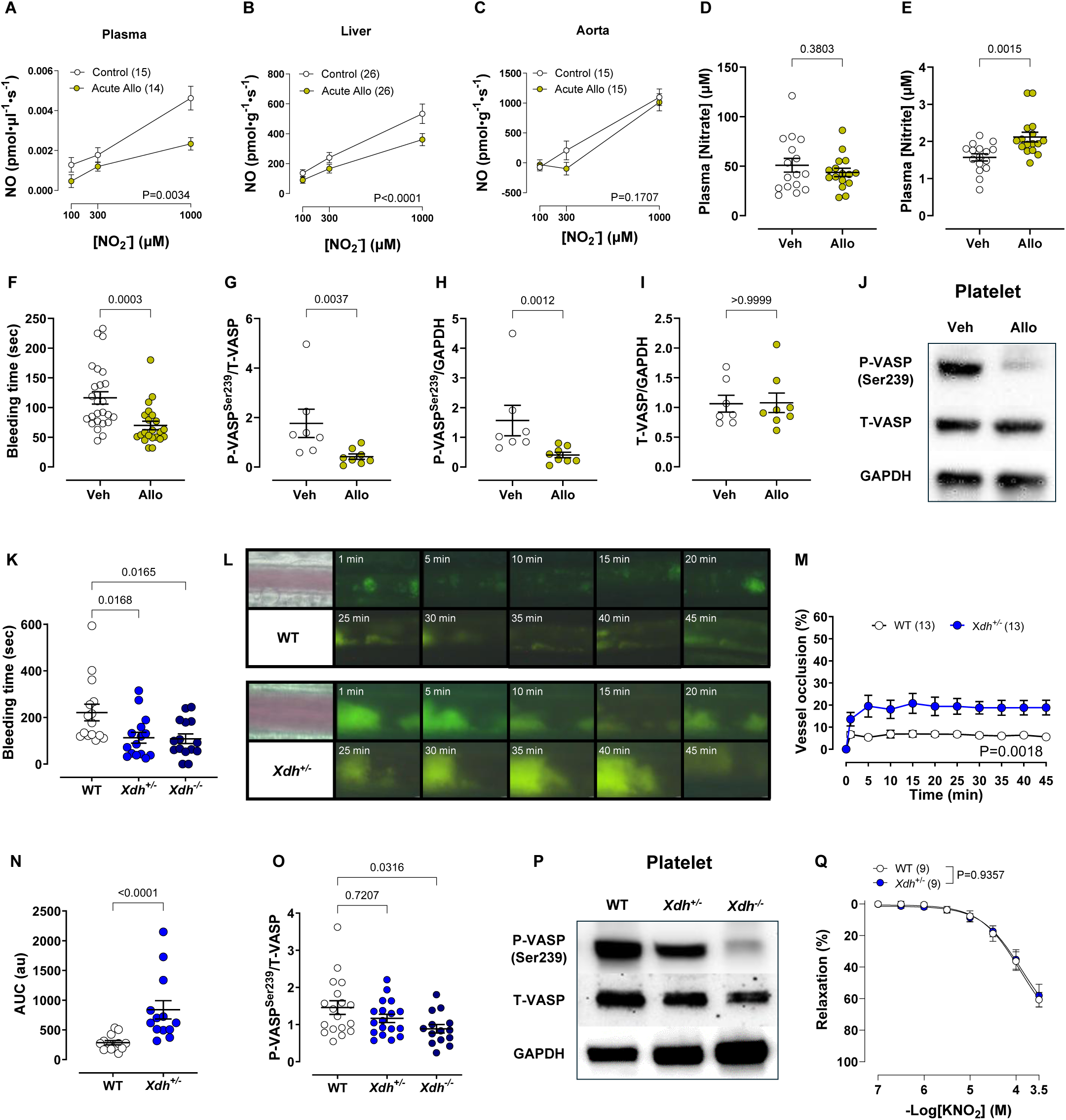
XOR drives antithrombotic nitrate reduction *in vivo.* (**A**) Nitrite reductase activity at pH 6.8 in the plasma, (**B**) liver homogenates or (**C**) aortic tissue homogenates of mice in control conditions (control) or after 30 minutes of acute treatment with 100 μM allopurinol (Acute Allo). (**D**) Plasma nitrate and (**E**) nitrite concentrations in wild-type animals treated for two weeks with allopurinol (Allo, 1 mM allopurinol, containing 1.2 mM NaOH) or vehicle (Veh, 1.2 mM NaOH) in the drinking water. (**F**) Tail bleeding time. (**G**) Quantitative analyses of vasodilator-stimulated phosphoprotein (VASP) phosphorylation at serine 239 (P-VASP^Ser239^) normalised to total VASP (T-VASP) expression in platelets, (**H**) P-VASP^Ser239^ normalised to GAPDH, and (**I**) T-VASP normalised to GAPDH. (**J**) Representative Western blot images of P-VASP^Ser239^, T-VASP and GAPDH. (**K**) Tail bleeding time. (**L**) Representative images of thrombus formation. (**M**) Vessel occlusion over time, represented as a percentage of thrombus formation to the total vessel area. (**N**) Area under the curve (AUC) of vessel occlusion over time. (**O**) Quantitative analyses of P-VASP^Ser239^ normalised to T-VASP expression in platelets. (**P**) Representative Western blot images of P-VASP^Ser239^, T-VASP and GAPDH. (**Q**) Potassium nitrite (KNO_2_)-induced relaxation of aortic rings precontracted with U46619 in wild-type (WT) and XOR-heterozygous (*Xdh^+/-^*) mice. Data are shown as mean ± SEM (shown on the individual graphs). Statistical significance was determined using mixed-effect analyses (**A**-**C**), Student’s t-test (**D**), Mann-Whitney test (**E-I** and **N**), one-way ANOVA followed by Kruskal-Wallis test (**K** and **O**) or two-way ANOVA (**M** and **Q**).

### Genetic XOR-deficiency promotes platelet hyperreactivity and thrombosis

To further establish a physiological role for XOR, haemostatic function was assessed in mice with genetic deletion of *Xdh*. XOR expression (Western blotting and flow cytometry) was absent from platelets across all the genotypes but readily detected in liver tissue, indicating that platelet NO signalling is regulated by extra-platelet sources of XOR (**Supplementary Figure 6C-F**). Despite normal platelet count and size (**Supplementary Figure 6G-I**) *Xdh^+/-^*mice exhibited a clear prothrombotic phenotype characterised by a shortened bleeding time (**Figure 3K**) and enhanced vascular injury response (**Figure 3L-N and Supplementary Figure 3J**), with no effects upon thrombus kinetics (**Supplementary Figure 6K&L**). These changes were accompanied by reduced VASP phosphorylation (**Figure 3 O&P and Supplementary Figure 6M-N**), indicating impaired NO-cGMP signalling. Vascular responses to nitrite and endothelium-dependent agonists remained largely preserved in heterozygous mice, suggesting that platelet regulation is more sensitive than vascular relaxation to reductions in XOR activity (**Figure 3Q and Supplementary Figure 6O&P**).

Mechanistically, platelets from *Xdh^+/-^* mice demonstrated enhanced collagen-induced aggregation and exaggerated calcium mobilisation in response to ionomycin, whereas P-selectin expression and platelet-leukocyte interactions were unaffected (**Figure 4**, **Supplementary Figure 6Q&R**). These observations identify dysregulated calcium handling as a likely downstream consequence of impaired XOR-dependent NO signalling and provide a cellular explanation for the prothrombotic phenotype.

**Figure 4:**
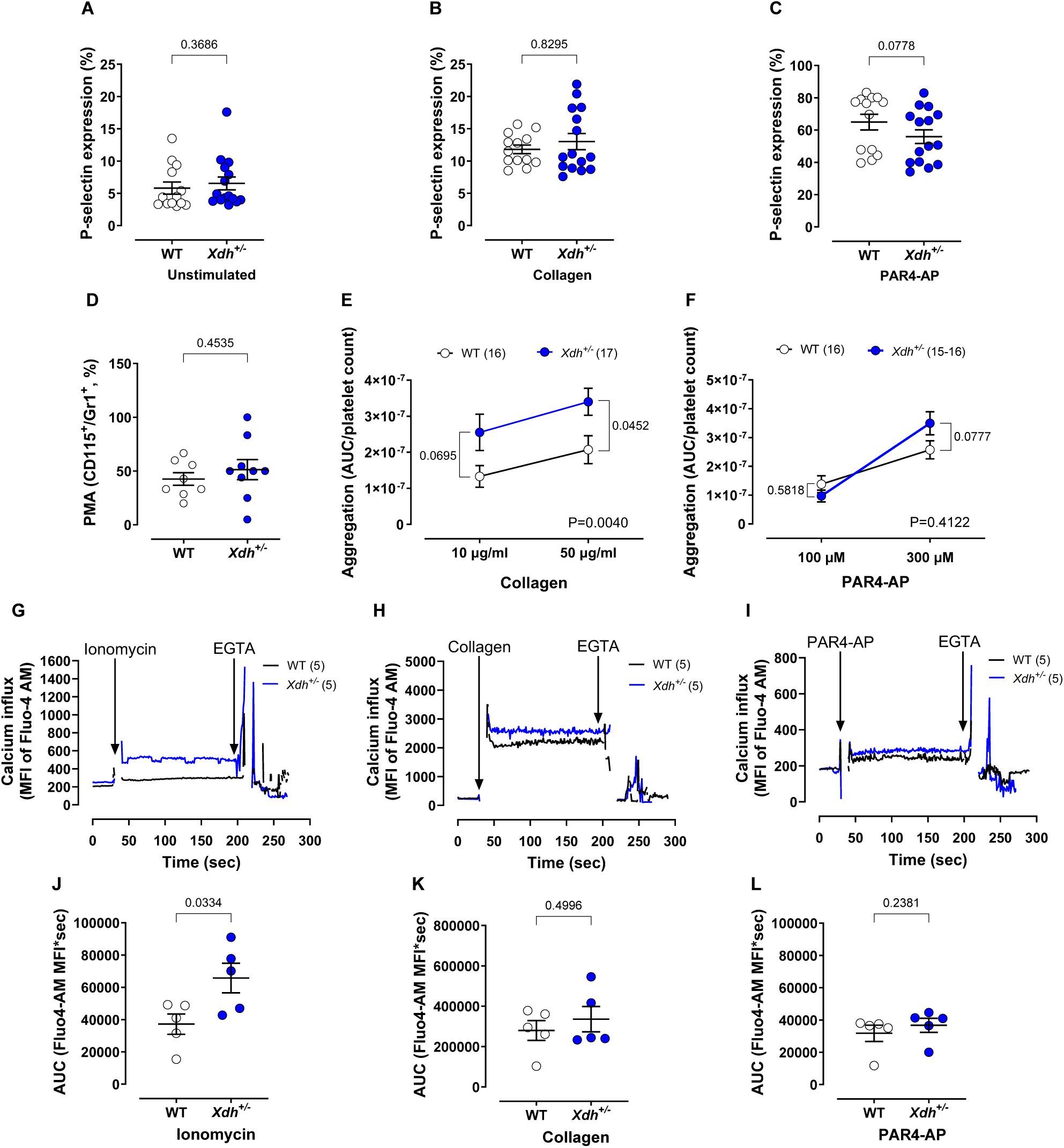
Modest effects of XOR deletion on *ex vivo* determined platelet activity. P-selectin expression on (**A**) unstimulated platelets, (**B**) stimulated with 50 μg/mL collagen or (**C**) stimulated with 300 μM PAR4-AP. (**D**) Inflammatory platelet-monocyte aggregates (PMA, CD115^+^/Gr1^+^). Platelet aggregation induced by collagen (**E**) and PAR4-AP (**F**), normalised to platelet count. Platelet calcium influx expressed as median fluorescence intensity (MFI) following activation with 10 μM ionomycin (**G**), 50 μg/mL collagen (**H**), or 300 μM PAR4-AP (**I**). Area under the curve (AUC) of the MFI during agonist activation (50 to 180 seconds period) with ionomycin (**J**), collagen (**K**) or PAR4-AP (**L**). Data are shown as mean ± SEM (shown on the individual graphs). Statistical significance was determined using Mann-Whitney test (**A-C** and **L**), Student’s t-test (**D, J** and **K**) and mixed-effect analysis followed by Šídák’s multiple comparisons test (**E** and **F**).

### The anti-thrombotic effects of dietary nitrate supplementation are absent in *Xdh^+/-^* mice

We next tested whether the beneficial anti-thrombotic effects of dietary nitrate are mediated by XOR. Although dietary nitrate supplementation increased plasma nitrate and nitrite levels similarly in both WT and in *Xdh^+/-^* mice (**Figure 5A&B**), only in WT animals was this accompanied by a prolonged bleeding time (**Figure 5C**) and reduced the thrombus formation (**Figure 5D-F**; (**Supplementary Figure 7A-C**). Consistent with impaired downstream signalling, platelet cGMP levels trended to be lower in the nitrate-treated *Xdh^+/-^* animals compared to WT mice (**Figure 5G**). As with the above observations, these changes were evident *in vivo* despite no alterations in *ex vivo* assessed standard markers of platelet activation (**Figure 5H-M; Supplementary Figure 7D&E**). These data demonstrate that increased substrate availability alone is insufficient to confer protection and establish XOR as an essential mediator linking dietary nitrate to platelet NO signalling and antithrombotic activity.

**Figure 5:**
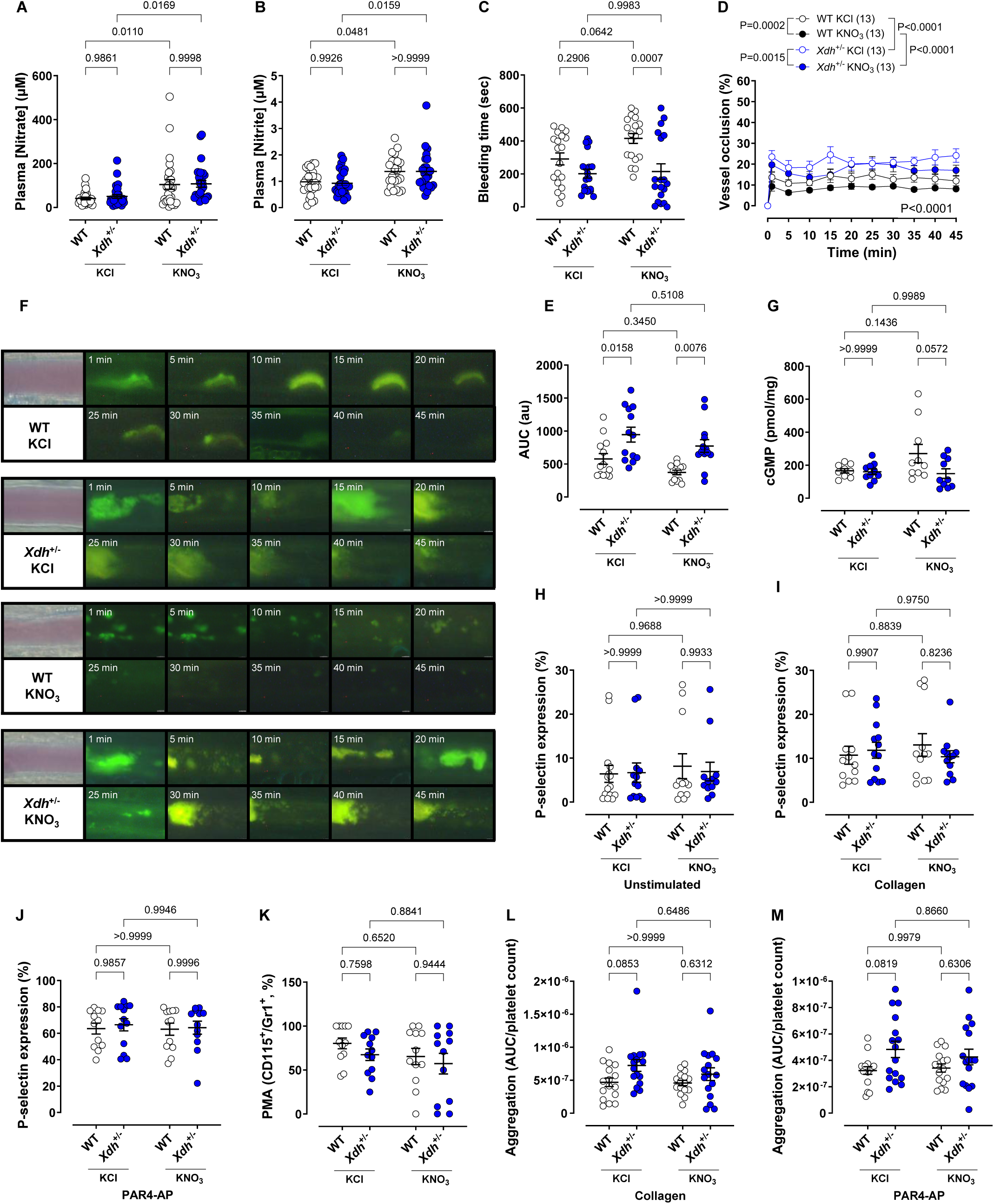
Global *Xdh* deletion eliminates antithrombotic nitrite reductase activity *in vivo*. (**A**) Plasma nitrate and (**B**) nitrite concentrations in wild-type (WT) and XOR-heterozygous (*Xdh^+/-^*) mice treated for two weeks with potassium nitrate (15 mM KNO_3_) in the drinking water or potassium chloride (15 mM KCl) as a control. (**C**) Tail bleeding time. (**D**) Vessel occlusion over time, represented as a percentage of thrombus formation to the total vessel area. (**E**) Area under the curve (AUC) of vessel occlusion over time. (**F**) Representative images of thrombus formation. (**G**) Platelet cGMP concentration. P-selectin expression on (**H**) unstimulated platelets, (**I**) stimulated with 50 μg/mL collagen or (**J**) 300 μM PAR4-AP. (**K**) Inflammatory platelet-monocyte aggregates (PMA, CD115^+^/Gr1^+^). (**L**) Platelet aggregation induced by 50 μg/mL collagen and (**M**) 300 μM PAR4-AP, normalised to platelet count. Data are shown as mean ± SEM (shown on the individual graphs). Statistical significance was determined using two-way ANOVA followed by Šídák’s multiple comparisons test.

### Hepatic XOR is the principal source of anti-thrombotic nitrite reductase activity

Since the liver is the major site of XOR generation, we generated mice lacking XOR selectively in hepatocytes (HXOR KO). Hepatic deletion markedly reduced hepatic XOR expression (**Figure 6A&B**) and impaired nitrite responses in the aorta, evidenced by blunted KNO_2_-induced vasorelaxation (**Figure 6C**), while endothelium-dependent (ACh) and direct NO-induced (Sper-NO) relaxation remained unchanged (**Supplementary Figure 8A&B**). Loss of liver-derived XOR was associated with reduced platelet VASP^Ser239^ phosphorylation (**Figure 6D&E and Supplementary Figure 8C&D**), increased platelet aggregation in response to collagen and PAR4-AP (**Figure 6F&G**) and shorter bleeding time (**Figure 6H-I, Supplementary Figure 8E&F**), closely recapitulating the phenotype observed in global XOR-deficient mice. These findings demonstrate that hepatic XOR is a key systemic source of nitrite reductase activity and plays a central role in maintaining platelet NO-cGMP signalling and thromboresistance.

**Figure 6:**
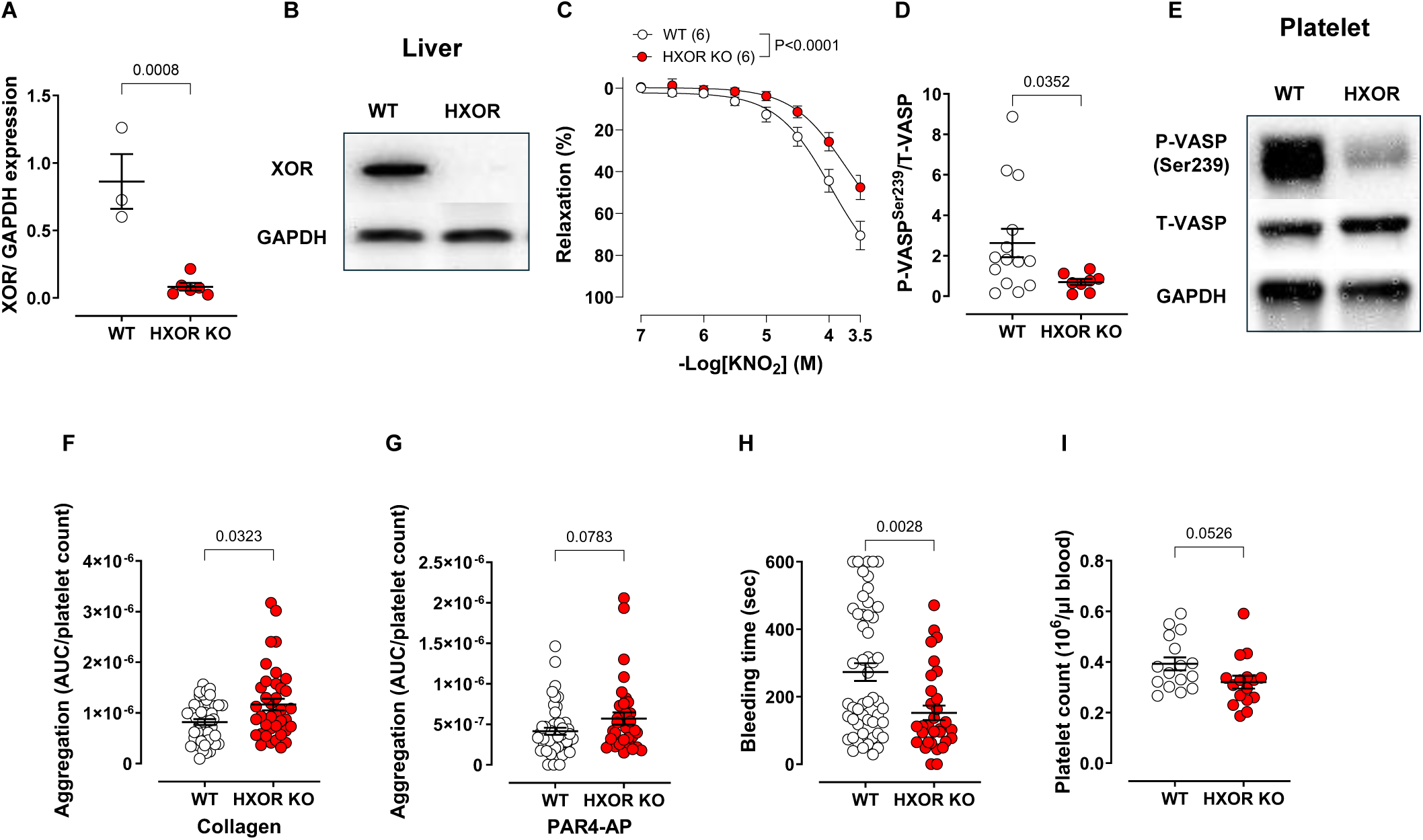
Hepatocyte-specific deletion of XOR recapitulates the prothrombotic phenotype of global *Xdh* deletion. (**A**) Quantitative analyses of XOR expression normalised to GAPDH in the liver of wild-type (WT) and hepatocyte-specific XOR-deficient (HXOR KO) mice. (**B**) Representative Western blot images of XOR and GAPDH. (**C**) Potassium nitrite (KNO_2_)-induced relaxation of aortic rings precontracted with phenylephrine in wild-type (WT) and HXOR KO mice. (**D**) Quantitative analyses of vasodilator-stimulated phosphoprotein (VASP) phosphorylation at serine 239 (P-VASP^Ser239^) normalised to total VASP (T-VASP) expression in platelets. (**E**) Representative Western blot images of P-VASP^Ser239^, T-VASP and GAPDH. Platelet aggregation induced by 50 μg/mL collagen (**F**) and 300 μM PAR4-AP (**G**). (**H**) Tail bleeding time. (**I**) Platelet count. Data are shown as mean ± SEM (shown on the individual graphs). Statistical significance was determined using Student’s t-test (**A** and **G**), two-way ANOVA (**C**) or Mann-Whitney test (**D**, **F**, **H** and **I**).

## Discussion

Our observations establish a new paradigm for platelet signalling, i.e **hepatic XOR is a critical source of nitrate-derived antithrombotic NO signalling**. The principal finding of this study is that XOR-dependent reduction of inorganic nitrite provides a tonic, physiologically active brake upon platelet activation and thrombosis *in vivo*, and that the liver is the dominant source of the reductase activity that sustains it. Four lines of evidence support this conclusion. First, dietary nitrate corrected the exaggerated thrombotic response of two mechanistically distinct models of endothelial dysfunction, the *eNOS*^-/-^ and the *ApoE*^-/-^ mouse, without altering the expression of XOR or indeed eNOS (at least in the *ApoE^-/-^* mice). Second, pharmacological inhibition of XOR with allopurinol suppressed nitrite reductase activity in plasma and liver, elevated circulating nitrite, shortened bleeding time and reduced platelet VASP phosphorylation at Ser239. Third, mice carrying a single functional *Xdh* allele were spontaneously prothrombotic and were entirely refractory to the antithrombotic actions of dietary nitrate despite equivalent elevations in plasma nitrate and nitrite. Fourth, hepatocyte-selective deletion of *Xdh* recapitulated the phenotype of global XOR deficiency. Since XOR protein was undetectable in platelets themselves, these observations together describe an inter-organ, endocrine-like axis in which hepatic XOR generates NO from circulating nitrite to restrain platelet reactivity at a distance.

That dietary nitrate restored haemostatic and thrombotic responses in *eNOS*^-/-^ mice is notable, since these animals lack the canonical endothelial source of platelet-directed NO and display enhanced haemostasis^17^. Our data extend this observation by showing that, although bleeding times were preserved at baseline, thrombus burden following FeCl_3_ injury was markedly increased, and that both endpoints were increased or normalised respectively by nitrate supplementation. Comparable protection was also observed in *ApoE*^-/-^ mice, where *in vivo* nitrate treatment improved relaxation not only to KNO_2_ but also to acetylcholine and spermine-NO and normalised phenylephrine-induced contraction. These findings are consistent with the anti-inflammatory and NO-sparing actions of inorganic nitrate described previously in this model^12,28^ and with reduced scavenging of NO by reactive oxygen species. Importantly, nitrate supplementation did not alter hepatic or vascular XOR expression, aortic eNOS expression or eNOS Ser1177 phosphorylation, indicating that the benefit derives from increased substrate delivery to a constitutively expressed reductase rather than from induction of the enzymatic machinery. These findings align with the antiplatelet effects of nitrate and nitrite reported in healthy volunteers^13–15^ and in murine models, and with the observation that *in vivo* platelet regulation depends critically upon NO but not upon eNOS itself^29,30^.

*In vitro* evidence strongly supports a role for XOR as a molybdopterin-dependent nitrite reductase, generating NO from nitrite at rates that increase as pH and pO_2_ decrease^31–33^. Whether this activity is quantitatively and functionally relevant *in vivo*, and whether it serves a discrete physiological function with respect to platelet function, has been much less certain. The present data addresses both questions. Chronic *in vivo* allopurinol treatment, to inhibit XOR activity, produced a pharmacodynamic profile explainable by blocked nitrite reductase activity of plasma nitrite accumulation whilst nitrate remained unchanged. This was accompanied by shortened bleeding time and reduced platelet VASP^Ser239^ phosphorylation (supporting previous evidence linking nitrite and VASP phosphorylation^34,35^), indicating a fall in cGMP-dependent signalling. Because allopurinol also lowers urate generation and has actions beyond XOR inhibition, we next sought genetic corroboration of these findings. *Xdh*^+/-^ mice reproduced the platelet phenotype in full, also expressing shortened bleeding time, enhanced thrombus formation, reduced platelet VASP phosphorylation and enhanced collagen-induced aggregation. These findings fit well with our recent observations of reduced platelet cGMP levels in both global and liver-selective *Xdh* knockout mice^16^. It is particularly noteworthy that a 50% reduction in gene dosage (i.e. heterozygotes) was sufficient to produce a prothrombotic phenotype, implying little functional reserve within the pathway; but this occurred whilst aortic relaxation to nitrite and to endothelium-dependent agonists essentially remained intact. Platelet function therefore appears to be a more sensitive reporter of XOR-derived NO than vascular smooth muscle relaxation, which may reflect the very low cGMP concentrations required to inhibit platelet activation^36^.

We were unable to detect XOR using selective antibodies by Western blotting or by flow cytometry of intracellular or surface epitopes, using either citrate or heparin as anticoagulant for blood collection. The latter is a particularly relevant control, since heparin displaces XOR bound to glycosaminoglycans^37–39^. XOR is synthesised predominantly in liver and intestine^40^, is released into the circulation, and binds avidly to sulphated glycosaminoglycans on the endothelial surface^38,41^, where it retains catalytic activity. Our hepatocyte-specific knockout provides direct *in vivo* evidence that this hepatic pool is the functionally dominant one. Indeed, HXOR KO mice showed markedly reduced hepatic XOR expression, blunted KNO_2_-induced aortic vasorelaxation, reduced platelet VASP^Ser239^ phosphorylation, enhanced aggregation to collagen and PAR4-AP, and shortened bleeding time, thus closely phenocopying global XOR deficiency. Critically, relaxation to acetylcholine and spermine-NO was unaffected, demonstrating an insufficiency in nitrite bioactivation rather than a generalised loss of endothelial function or smooth muscle NO responsiveness. These findings position the liver as an endocrine organ for NO generation, complementing the erythrocytic XOR-dependent nitrite reduction we have previously described in humans, including in hypertension^11,42^ and the age-dependent decline in XOR nitrite reductase activity recently reported^16^.

These data also resolve an apparent inconsistency in our own results. Nitrite reductase activity was evidenced in aortic homogenates but was not suppressed by allopurinol, yet hepatocyte-specific XOR deletion blunted the aortic response to nitrite. An interpretation of these results is that the activity measured in vessel wall homogenates is substantially attributable to non-XOR reductases, potentially including aldehyde oxidase and haem proteins, together with acidic disproportionation^43,44^. In contrast, the functionally relevant reductase acting in the intact circulation is delivered from the liver as a circulating enzyme that then is bound to the endothelial surface. Homogenate assays do not permit discrimination of this type of compartmentalisation.

Downstream of NO, the signalling read-outs observed were internally consistent across all three interventions. P-VASP^Ser239^ is considered a largely selective substrate of protein kinase G^45^, and its phosphorylation decreased with allopurinol treatment, with global and hepatocyte-specific deletion, and platelet cGMP trended lower in nitrate-treated *Xdh*^+/-^ animals compared with wild-type. Loss of cGMP-dependent signalling is well established to promote platelet adhesion and aggregation^46–48^. Our work interrogating platelet calcium levels offers a potential cellular explanation for the phenotype. Platelets from *Xdh*^+/-^ mice showed exaggerated calcium mobilisation in response to ionomycin, whereas responses to collagen and PAR4-AP were unaltered, suggesting that the impairment likely occurs downstream of receptor signalling, in the PKG-dependent steps that limit and clear cytosolic calcium. Consistent with this possibility are the findings that P-selectin expression and platelet-leukocyte interactions, which indicate granule secretion rather than calcium handling per se, were unaffected.

A recurring feature of our findings is the dissociation between *in vivo* phenotype and conventional *ex vivo* markers of platelet activation. Across every model, bleeding time and thrombus formation responded robustly while P-selectin expression, platelet-monocyte aggregates and impedance aggregometry were largely unchanged. We suggest two potential explanations. Mechanistically, XOR-catalysed nitrite reduction is strongly favoured by the hypoxia^49^ and probable acidosis that characterise a growing thrombus, conditions not present in oxygenated, buffered *ex vivo* assays. Secondly, since the reductase is extrinsic to the platelet, isolating platelets from the vascular compartment removes the NO source itself. This is consistent with evidence that nitrite-mediated platelet inhibition requires erythrocytes and deoxygenation^34^, and that circulating nitrite and nitrate levels affect murine platelet function *in vivo*^50^. Methodologically, the *in vivo* assays also allow integration of an NO signal on thrombus formation over a 45-minute period and thus may simply be more sensitive. Notably, nitrate treatment altered the magnitude of thrombus formation without affecting the kinetics of thrombus growth or its stability, indicating an effect on platelet recruitment rather than an alteration in thrombus structure. The broader implication is cautionary: *ex vivo* aggregometry may systematically underestimate the antiplatelet efficacy of nitrate- and nitrite-based interventions, and *in vivo* endpoints should be preferred when evaluating their potential.

We propose two principal translational implications of our findings. The first is that the nitrate-nitrite-NO pathway continues to operate, and is therefore therapeutically exploitable, where endothelial-NO-derived generation has failed: a setting in which antithrombotic protection is most needed^10^. The second relates to XOR inhibitor prescription. XOR inhibitors are often prescribed long term to very large numbers of patients with gout and whose cardiovascular risk profile is elevated. Our data show that XOR inhibition is not neutral with respect to platelet function: it removes a tonic antithrombotic NO signal and abolishes the benefit of dietary but also endogenously generated inorganic nitrate. We speculate that this effect may have contributed to the neutral outcome of the ALL-HEART trial, in which allopurinol failed to improve cardiovascular outcomes in patients with ischaemic heart disease^51^, despite earlier positive mechanistic studies in stable angina^52^ and cardiovascular safety data in gout^53^ (at least of febuxostat over allopurinol). Such an effect could be readily tested by assessing the vascular and antiplatelet benefits of a nitrate-rich diet in patients receiving allopurinol or febuxostat. Of course the benefits of not inhibiting XOR and allowing efficacy of nitrate has to be balanced against the oxidant-generating capacity of XOR, which is a major contributing rationale for inhibiting it^54,55^. The net effect of XOR inhibition is likely to be context-dependent, and our data argue that the loss of nitrite reductase function needs to be considered as part of that balance.

Several limitations should be acknowledged. Global XOR deletion causes early postnatal lethality, typically within four weeks, owing to the accumulation of hypoxanthine and xanthine crystals in the kidney^19,56^; most experiments were therefore performed in heterozygous (*Xdh*^+/-^) animals, with global knockout mice used only in a limited number of studies, and we cannot formally establish whether complete loss of XOR produces a proportionally greater phenotype, although the hepatocyte-specific data suggest that it does. Animals were studied at 4-8 weeks of age, so the *ApoE*^-/-^ data reflect early rather than advanced atherosclerosis. The FeCl_3_ model is an oxidant-driven injury that does not fully recapitulate plaque rupture, although the concordance between this assay and tail bleeding across every intervention argues against a model-specific artefact. Differences in platelet cGMP reached only a trend, reflecting the limited yield of murine platelets, and VASP^Ser239^ phosphorylation was therefore used as the principal proxy for pathway activity. Hepatocyte-specific deletion does not exclude contributions from intestinal, endothelial, myeloid or erythrocyte-associated XOR, and the relative contribution of these pools may differ between mouse and human. Finally, although all studies were sex-balanced, they were not powered to detect sex differences; given the reported influence of sex on the antiplatelet response to dietary nitrate in humans^57^, this warrants further investigation.

In conclusion, we identify hepatic XOR as the principal source of the nitrite reductase activity that sustains platelet NO-cGMP signalling and thromboresistance *in vivo*. We also show that dietary nitrate acts through this pathway to restore antithrombotic protection when endothelial NO generation fails. These findings further substantiate the reposition of XOR from a purely deleterious oxidase and purine processor towards a protective component of NO homeostasis, define an unanticipated hepatic contribution to the regulation of platelet function, and raise the possibility that pharmacological inhibition of XOR carries a previously unrecognised prothrombotic cost.

## Acknowledgements

We thank Professor Tim D Warner for his technical advice and guidance regarding platelet *in vitro* assays.

## CreDit Nomenclature

TP: data collection, data curation, formal analysis, data interpretation, writing – original draft and takes responsibility for the data.

CPT: data collection, data curation, formal analysis, data interpretation, writing – original draft and takes responsibility for the data.

FF: data collection, data curation, formal analysis, data interpretation, and takes responsibility for the data, reviewing–original draft.

ND: data collection, reviewing – original draft.

HA: Methodology, reviewing – original draft.

NC: data collection, reviewing – original draft.

RSK: data collection, data curation, formal analysis, data interpretation, writing – original draft.

GM: data collection, reviewing – original draft.

MC: funding acquisition, investigation, reviewing – original draft.

AA: conceptualisation, funding acquisition, investigation, methodology, data interpretation, writing – original draft and takes responsibility for the data.

---

**Supplementary Figure 1:**
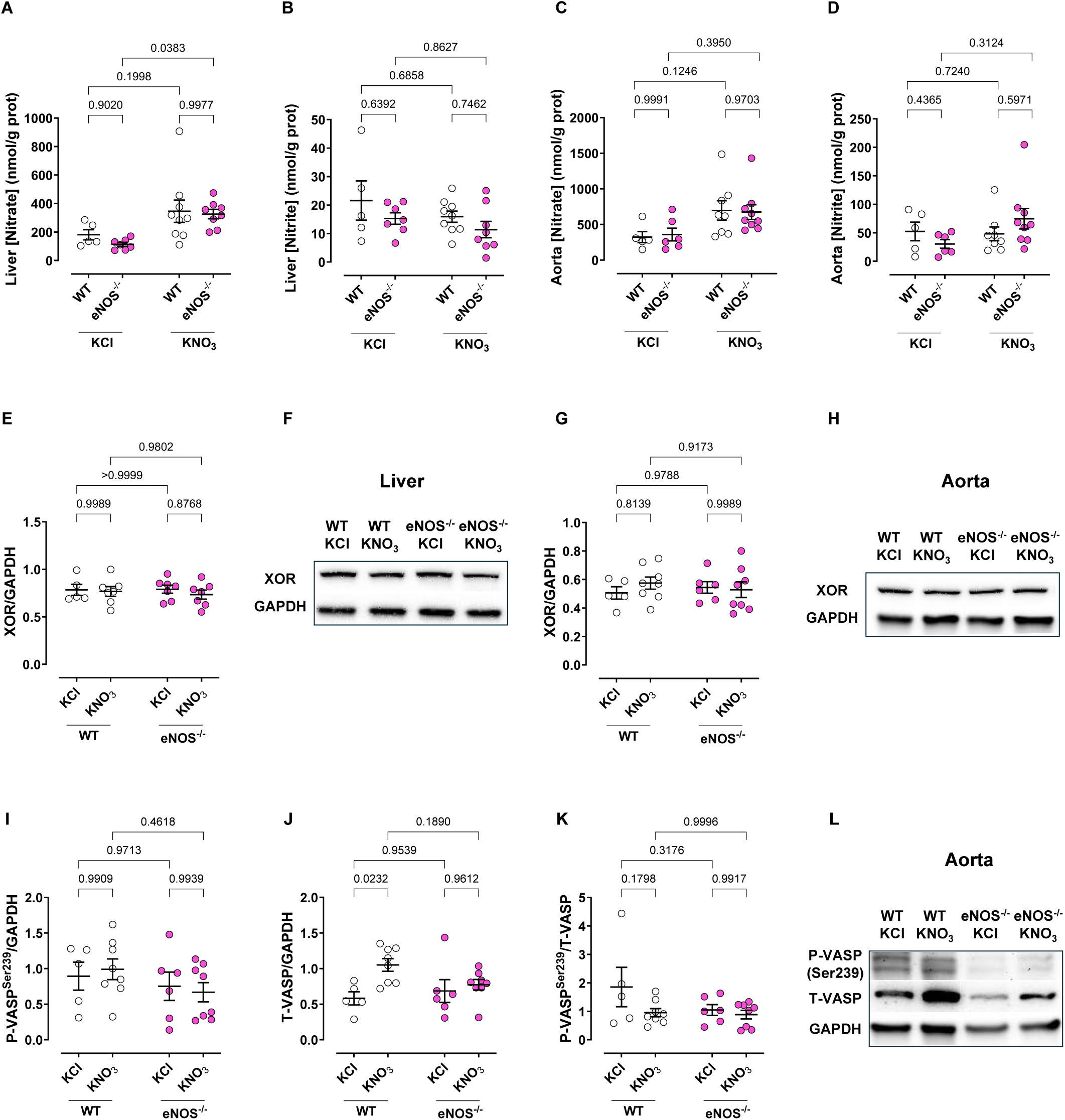
Nitrate (**A** and **C**) and nitrite (**B** and **D**) concentrations in liver (**A** and **B**) and aorta homogenates (**C** and **D**) from wild-type (WT) or endothelial nitric oxide synthase-deficient (eNOS^-/-^) mice treated for two weeks with potassium nitrate (15 mM KNO_3_) in the drinking water or potassium chloride (15 mM KCl) as a control. Quantitative analyses of XOR expression normalised GAPDH (**E** and **G**) and representative images (**F** and **H**) from liver (**E** and **F**) and aorta (**G** and **H**) homogenates. (**I**) Quantitative analyses of vasodilator-stimulated phosphoprotein (VASP) phosphorylation at serine 239 (P-VASP^Ser239^) normalised to GAPDH in the aorta. (**I**) Quantitative analyses of total VASP (T-VASP) normalised to GAPDH in the aorta. (**K**) Quantitative analyses of P-VASP normalised to T-VASP in the aorta. Representative Western blot images of P-VASP, T-VASP and GAPDH in the aorta. Data are shown as mean ± SEM (shown on the individual graphs). Statistical significance was determined using two-way ANOVA followed by Šídák’s multiple comparisons test.

**Supplementary Figure 2:**
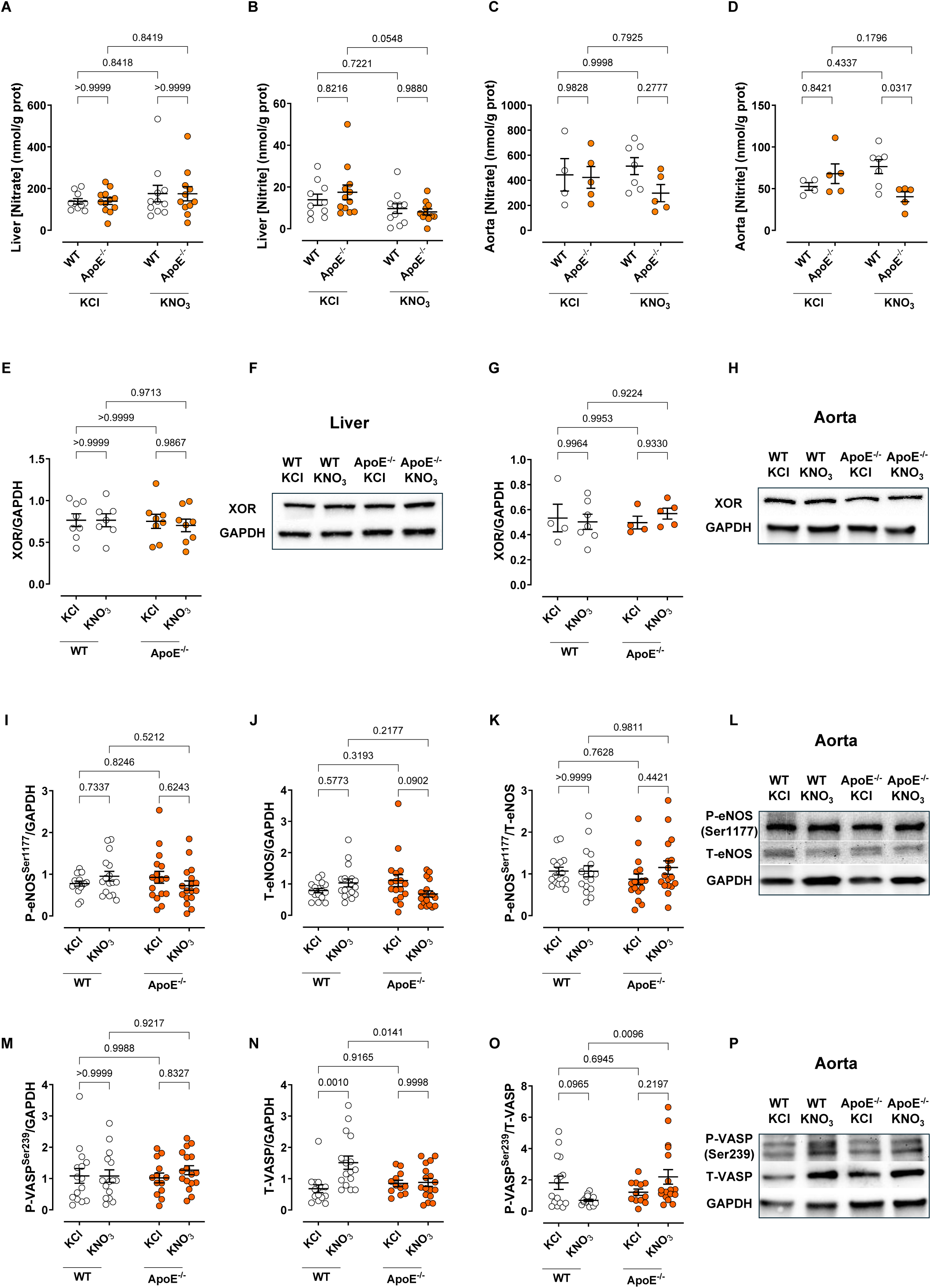
Nitrate (**A** and **C**) and nitrite (**B** and **D**) concentrations in liver (**A** and **B**) and aorta homogenates (**C** and **D**) from wild-type (WT) and apolipoprotein E-deficient (ApoE^-/-^) mice treated for two weeks with potassium nitrate (15 mM KNO_3_) in the drinking water or potassium chloride (15 mM KCl) as a control. Quantitative analyses of XOR expression normalised to GAPDH (**E** and **G**) and representative images (**F** and **H**) from liver (**E** and **F**) and aorta (**G** and **H**) homogenates. (**I**) Quantitative analyses of endothelial nitric oxide synthase phosphorylation at serine 1177 (P-eNOS^Ser1177^) normalised to GAPDH in the aorta. (**J**) Quantitative analyses of total eNOS (T-eNOS) normalised to GAPDH in the aorta. (**K**) Quantitative analyses of P-eNOS^Ser1177^ normalised to T-eNOS in the aorta. (**L**) Representative Western blot images of P-eNOS^Ser1177^, T-eNOS and GAPDH in the aorta. (**M**) Quantitative analyses of vasodilator-stimulated phosphoprotein (VASP) phosphorylation at serine 239 (P-VASP^Ser239^) normalised to GAPDH in the aorta. (**N**) Quantitative analyses of total VASP (T-VASP) normalised to GAPDH in the aorta. (**O**) Quantitative analyses of P-VASP normalised to T-VASP in the aorta. (**P**) Representative Western blot images of P-VASP, T-VASP and GAPDH in the aorta. Data are shown as mean ± SEM (shown on the individual graphs). Statistical significance was determined using two-way ANOVA followed by Šídák’s multiple comparisons test.

**Supplementary Figure 3:**
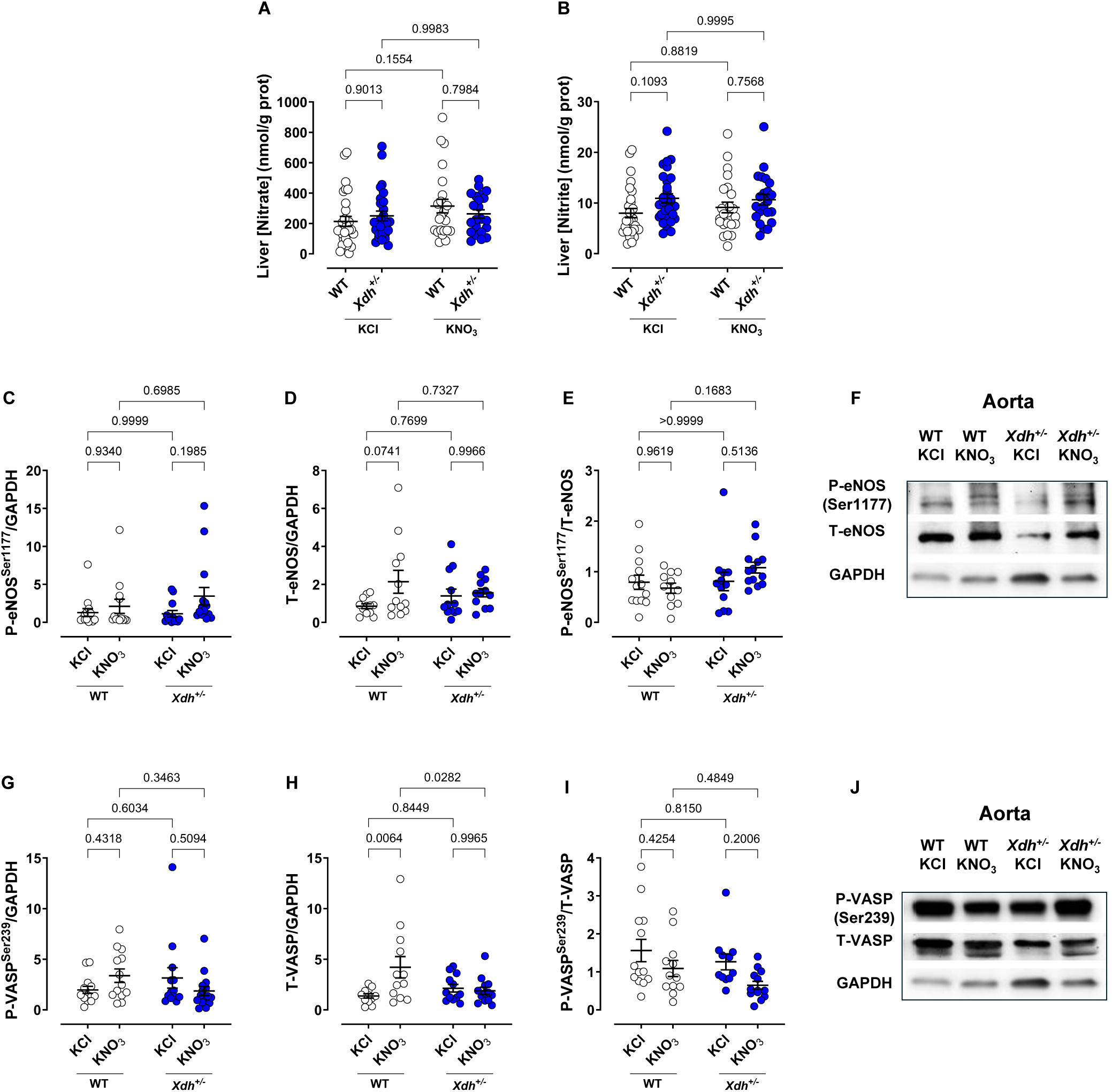
Nitrate (**A**) and nitrite (**B**) concentrations in liver from wild-type (WT) and XOR-heterozygous (*Xdh^+/-^*) mice treated for two weeks with potassium nitrate (15 mM KNO_3_) in the drinking water or potassium chloride (15 mM KCl) as a control. (**C**) Quantitative analyses of endothelial nitric oxide synthase phosphorylation at serine 1177 (P-eNOS^Ser1177^) normalised to GAPDH in the aorta. (**D**) Quantitative analyses of total eNOS (T-eNOS) normalised to GAPDH in the aorta. (**E**) Quantitative analyses of P-eNOS^Ser1177^ normalised to T-eNOS in the aorta. (**F**) Representative Western blot images of P-eNOS^Ser1177^, T-eNOS and GAPDH in the aorta. (**G**) Quantitative analyses of vasodilator-stimulated phosphoprotein (VASP) phosphorylation at serine 239 (P-VASP^Ser239^) normalised to GAPDH in the aorta. (**H**) Quantitative analyses of total VASP (T-VASP) normalised to GAPDH in the aorta. (**I**) Quantitative analyses of P-VASP normalised to T-VASP in the aorta. (**J**) Representative Western blot images of P-VASP, T-VASP and GAPDH in the aorta. Data are shown as mean ± SEM (shown on the individual graphs). Statistical significance was determined using two-way ANOVA followed by Šídák’s multiple comparisons test.

**Supplementary Figure 4:**
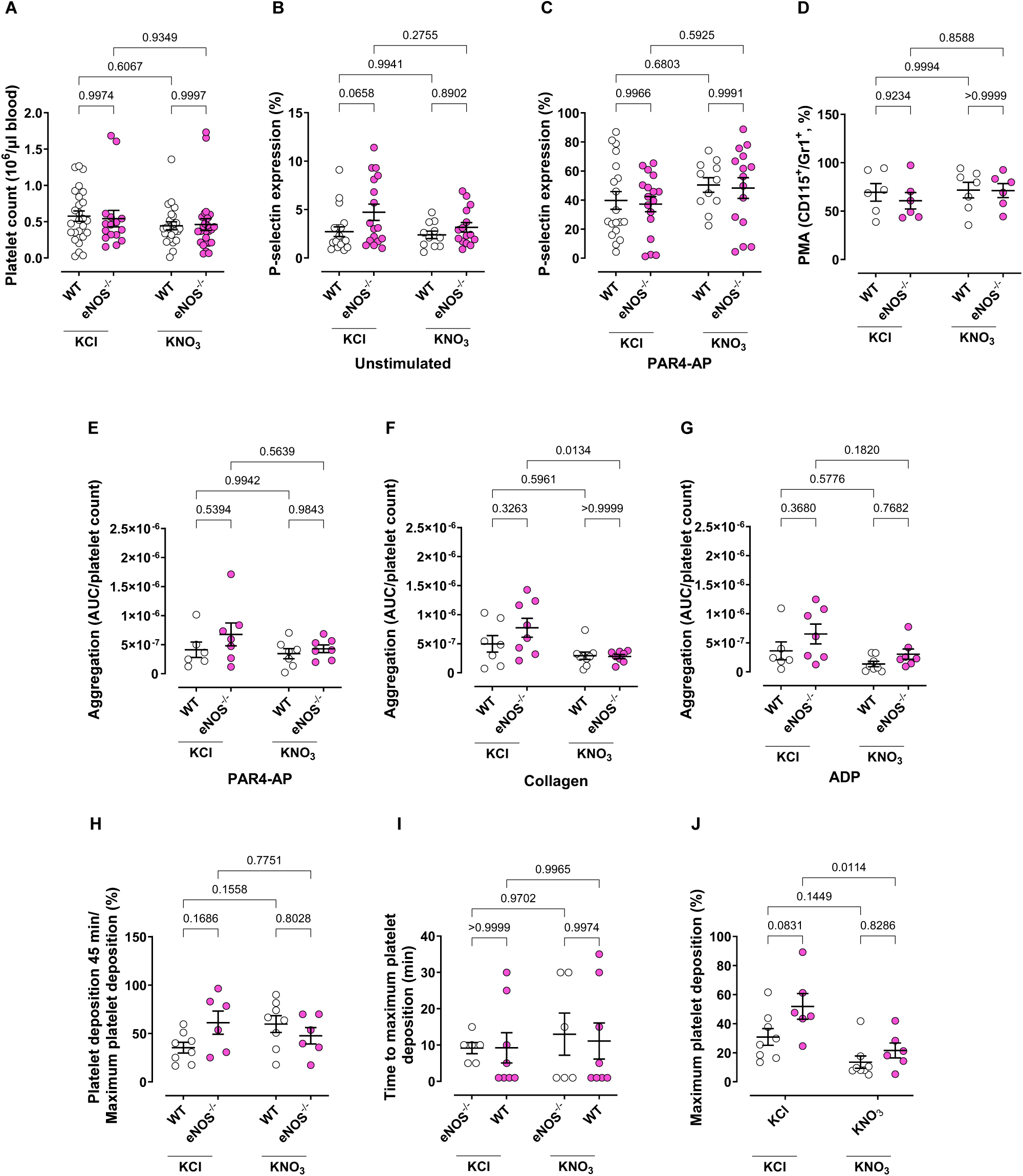
(**A**) Platelet count in wild-type (WT) endothelial nitric oxide synthase-deficient (eNOS^-/-^) mice treated for two weeks with potassium nitrate (15 mM KNO_3_) in the drinking water or potassium chloride (15 mM KCl) as a control. P-selectin expression on (**B**) unstimulated platelets or (**C**) platelets stimulated with 300 μM PAR4-AP. (**D**) Inflammatory platelet-monocyte aggregates (PMA, CD115^+^/Gr1^+^). Platelet aggregation induced by 300 μM PAR4-AP (**E**), 30 μM ADP (**F**), and 50 μg/mL collagen (**G**). (**H**) Platelet deposition at 45 minutes relative to maximum platelet deposition, (**I**) time to maximum platelet deposition, and (**J**) maximum platelet deposition following FeCl_3_ stimulation of thrombus formation. Data are shown as mean ± SEM (shown on the individual graphs). Statistical significance was determined using two-way ANOVA followed by Šídák’s multiple comparisons test.

**Supplementary Figure 5:**
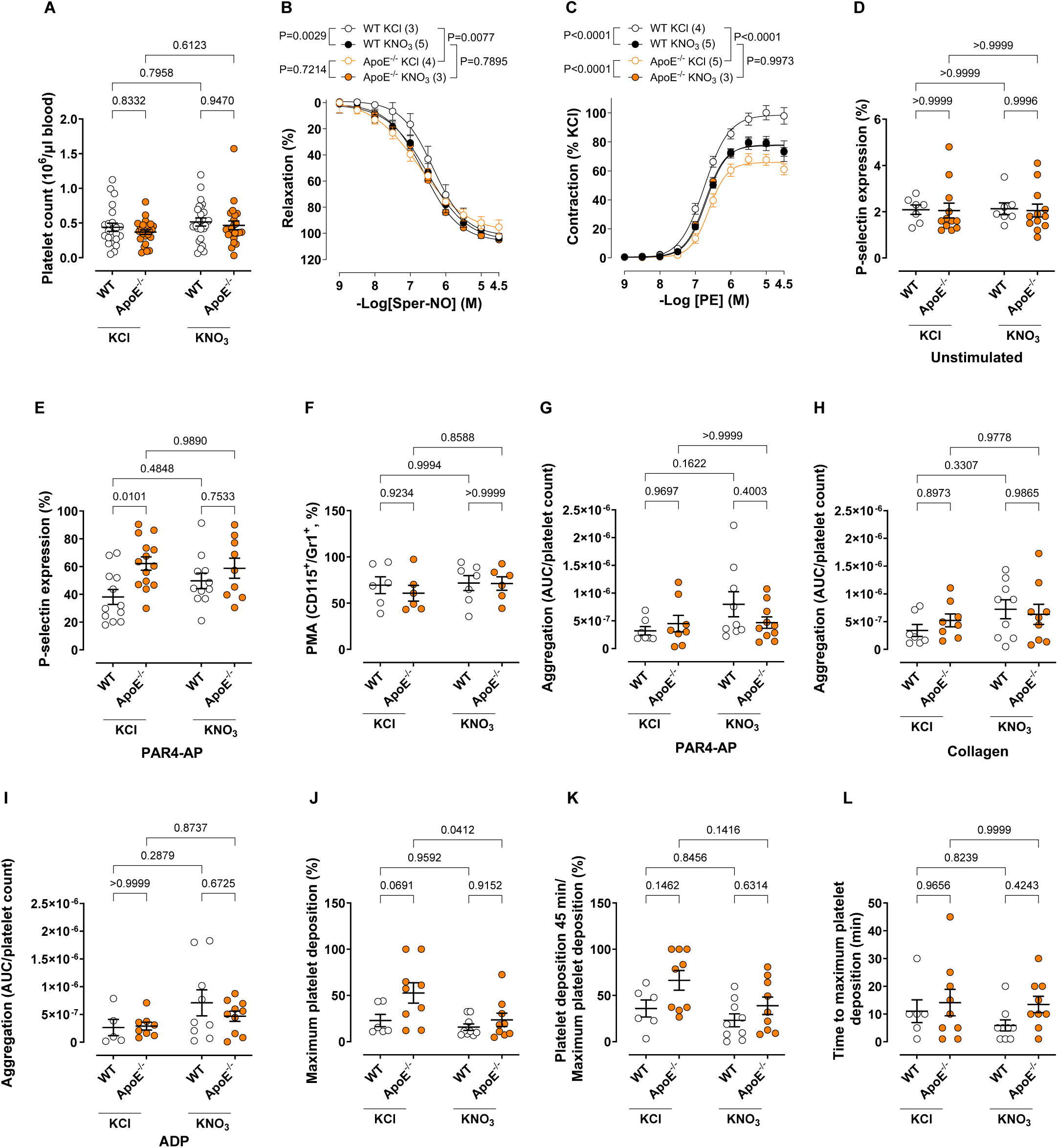
(**A**) Platelet count in wild-type (WT) and apolipoprotein E-deficient (ApoE^-/-^) mice treated for two weeks with potassium nitrate (15 mM KNO_3_) in the drinking water or potassium chloride (15 mM KCl) as a control. (**B**) Spermine-NO (Sper-NO)-induced relaxation of aortic rings precontracted with phenylephrine. (**C**) Phenylephrine (PE)-induced contraction of aortic rings. P-selectin expression on unstimulated platelets (**D**) or platelets stimulated with 300 μM PAR4-AP (**E**). (**F**) Inflammatory platelet-monocyte aggregates (PMA, CD115^+^/Gr1^+^). Platelet aggregation induced by 300 μM PAR4-AP (**G**), 50 μg/mL collagen (**H**), and 30 μM ADP (**I**). (**J**) Maximum platelet deposition, (**K**) platelet deposition at 45 minutes relative to maximum platelet deposition and (**L**) time to maximum platelet deposition following FeCl_3_ stimulation of thrombus formation. Data are shown as mean ± SEM (shown on the individual graphs). Statistical significance was determined using two-way ANOVA followed by Šídák’s multiple comparisons test.

**Supplementary Figure 6:**
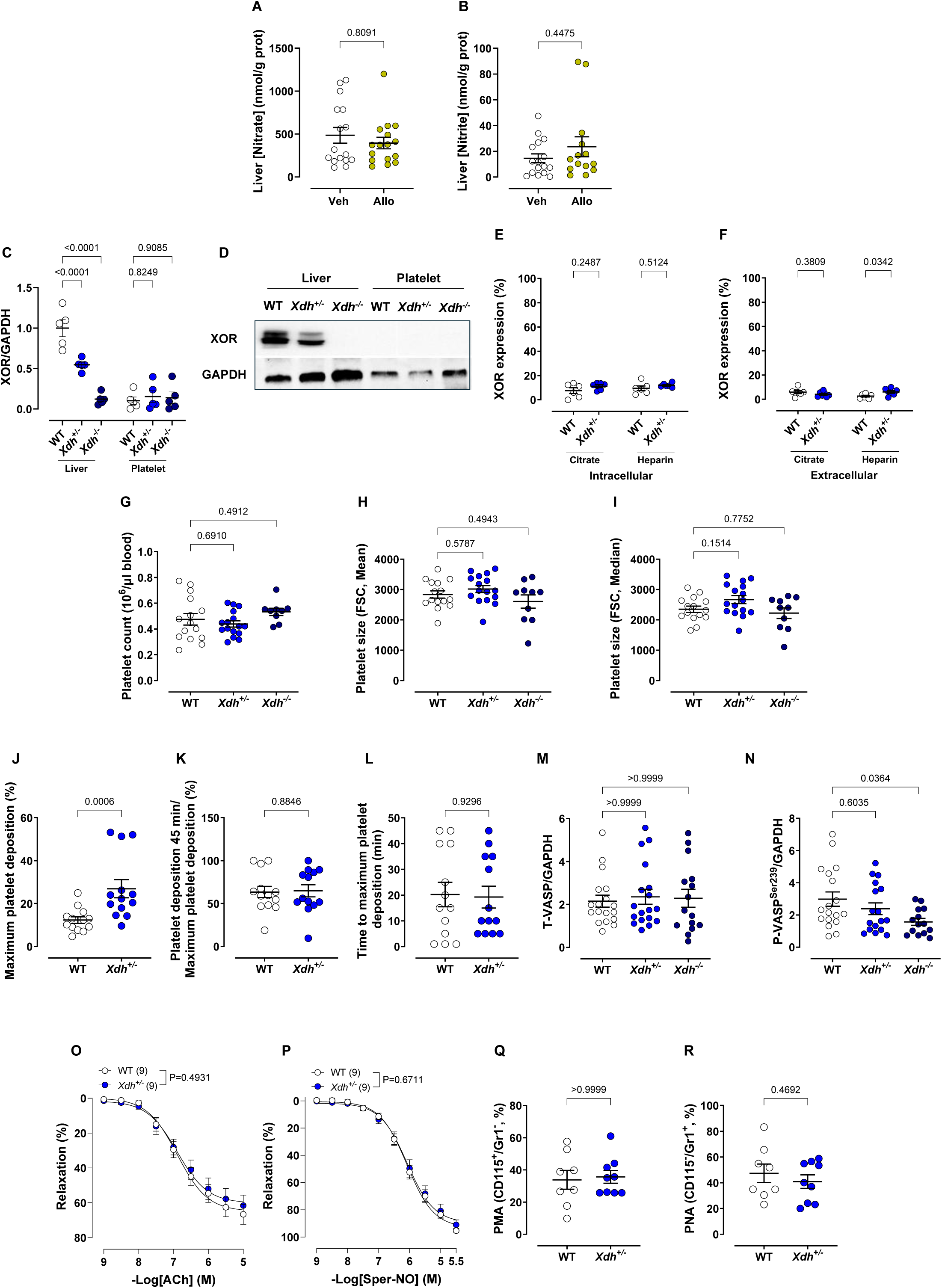
(**A**) Nitrate and (**B**) nitrite concentrations in liver homogenates from wild-type animals treated for two weeks with allopurinol (Allo, 1 mM allopurinol, containing 1.2 mM NaOH) or vehicle (Veh, 1.2 mM NaOH) in the drinking water. (**C**) Quantitative analyses of XOR expression normalised to GAPDH in the liver and platelets. (**D**) Representative Western blot images of XOR and GAPDH. Flow cytometry evaluation of intracellular (**E**) and extracellular (**F**) XOR expression in platelets isolated using either citrate or heparin as anticoagulant during blood collection. (**G**) Platelet count. Mean (**H**) and median (**I**) platelet size estimated by flow cytometry. (**J**) Maximum platelet deposition, (**K**) platelet deposition at 45 minutes relative to maximum platelet deposition, and (**L**) time to maximum platelet deposition following FeCl_3_ stimulation of thrombus formation. (**M**) Quantitative analyses of total vasodilator-stimulated phosphoprotein (T-VASP) expression and (**N**) VASP phosphorylation at serine 239 (P-VASP^Ser239^) expression, normalised to GAPDH expression in platelets. (**O**) Acetylcholine (ACh)- and (**P**) spermine-NO (Sper-NO)-induced relaxation of aortic rings precontracted with U19946 in wild-type (WT) and XOR-heterozygous (*Xdh^+/-^*) mice. (**Q**) Resident platelet-monocyte aggregates (PMA, CD115^+^/Gr1^-^). (**R**) Platelet-neutrophil aggregates (PNA, CD115^-^/Gr1^+^). Data are shown as mean ± SEM (shown on the individual graphs). Statistical significance was determined using Mann-Whitney test (**A, B, J, L** and **Q**), two-way ANOVA followed by Šídák’s multiple comparisons test (**C, E** and **F**), one-way ANOVA followed by Šídák’s multiple comparisons test (**G**) or Kruskal-Wallis test (**H, I, M** and **N**), two-way ANOVA (**O** and **P**) or Student’s t-test (**K** and **R**).

**Supplementary Figure 7:**
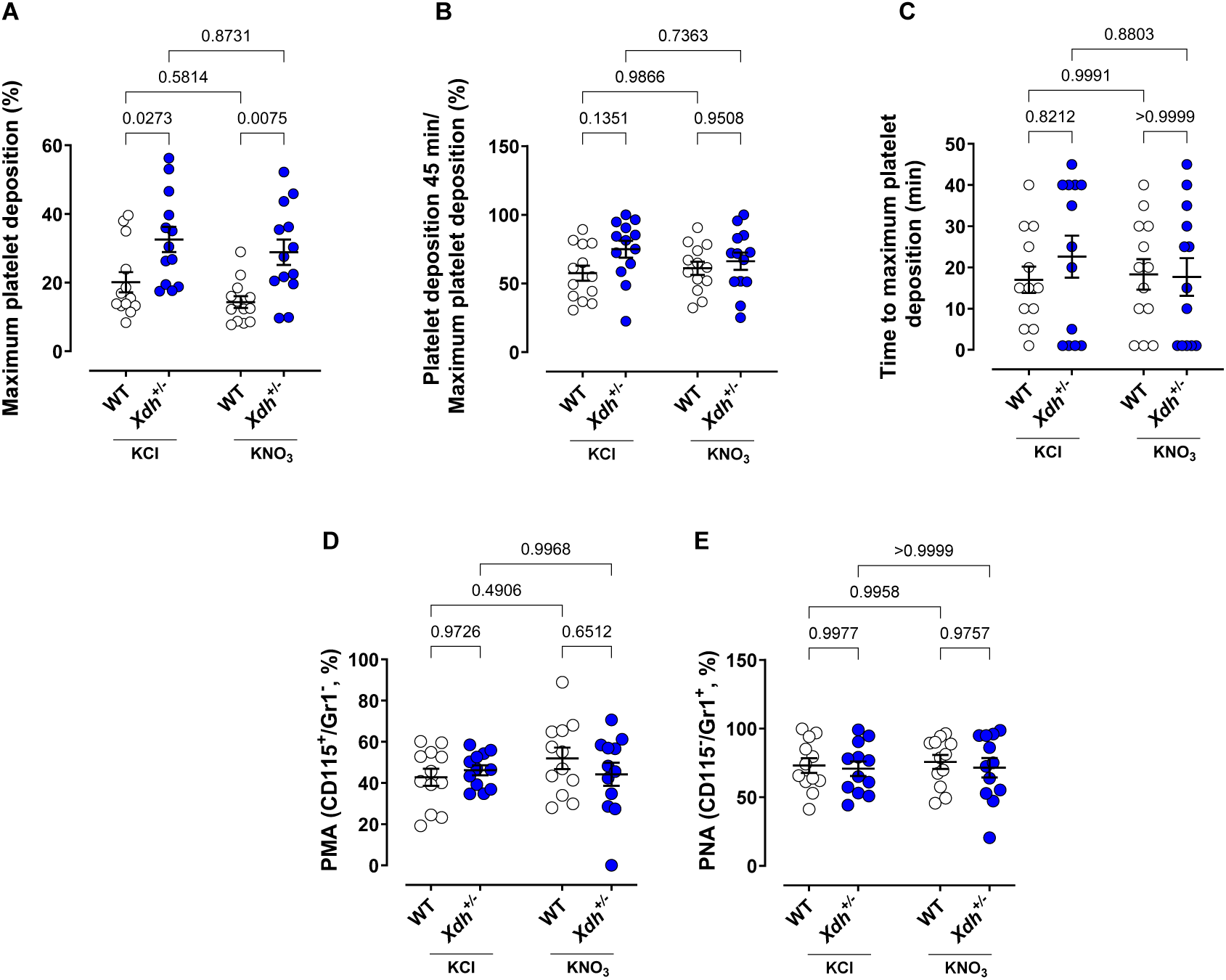
(**A**) Maximum platelet deposition, (**B**) platelet deposition at 45 minutes relative to maximum platelet deposition and (**C**) time to maximum platelet deposition following FeCl_3_ stimulation of thrombus formation from wild-type (WT) and XOR-heterozygous (*Xdh^+/-^*) mice treated for two weeks with potassium nitrate (15 mM KNO_3_) in the drinking water or potassium chloride (15 mM KCl) as a control. (**D**) Resident platelet-monocyte aggregates (PMA, CD115^+^/Gr1^-^). (**E**) Platelet-neutrophil aggregates (PNA, CD115^-^/Gr1^+^). Data are shown as mean ± SEM (shown on the individual graphs). Statistical significance was determined using two-way ANOVA followed by Šídák’s multiple comparisons test.

**Supplementary Figure 8:**
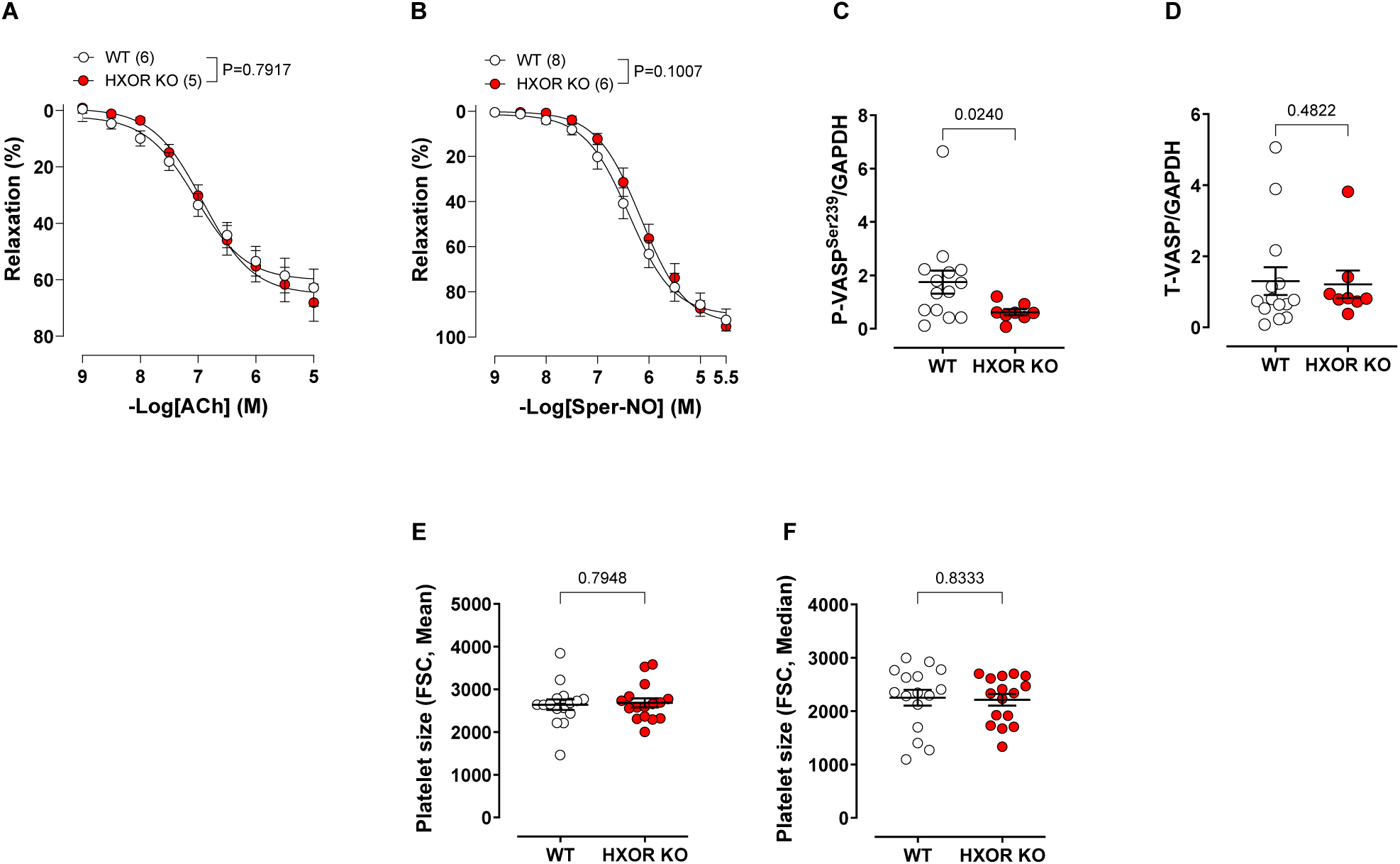
(**A**) Acetylcholine (ACh)- and (**B**) spermine-NO (Sper-NO)-induced relaxation of aortic rings precontracted with U19946 in wild-type (WT) and hepatocyte-specific XOR-deficient (HXOR KO) mice. (**C**) Quantitative analyses of vasodilator-stimulated phosphoprotein (VASP) phosphorylation at serine 239 (P-VASP^Ser239^) normalised to GAPDH in platelets. (**D**) Quantitative analyses of total VASP (T-VASP) expression normalised to GAPDH in platelets. Mean (**E**) and median (**F**) platelet size estimated by flow cytometry. Data are shown as mean ± SEM (shown on the individual graphs). Statistical significance was determined using two-way ANOVA (**A** and **B**), Student’s t-test (**C**, **E** and **F**) or Mann-Whitney test (**C**).

